# Ultrasound Directly Activates Sparse Neurons and Modulates Visual Circuits in Deafened Mice

**DOI:** 10.1101/2025.03.09.642207

**Authors:** Jiaru He, Jiejun Zhu, Xinxin Wang, Zihao Chen, Jin Yang, Zhen Yuan, Hongzhi Xu, Lei Sun, Zhihai Qiu

**Affiliations:** Guangdong Institute of Intelligence Science and Technology, Hengqin, Zhuhai, Guangdong 519031, China; Faculty of Health Sciences/Centre for Cognitive and Brain Sciences, University of Macau, Maca SAR, China; Department of Biomedical Engineering, The Hong Kong Polytechnic University, Hung Hom, Hong Kong SAR, P. R. China, 999077; Department of Neurosurgery, Huashan Hospital, Shanghai Medical College, Fudan University, 200040,China

**Keywords:** Ultrasound neuromodulation, Visual stimulation, Two-photon imaging, Auditory confound

## Abstract

Focused ultrasound neuromodulation (FUN) is widely regarded as a next-generation technology for neural modulation, with applications spanning rodents, non-human primates, and humans. While mechanistic studies are advancing, a persistent confound—auditory interference—casts doubt on whether ultrasound exerts direct mechanical effects or merely indirect auditory responses. To resolve this, we engineered a circular ultrasound transducer compatible with two-photon calcium imaging and examined its effects on the primary visual cortex (V1) of surgically deafened mice, eliminating auditory contributions. Our results reveal that ultrasound directly activates a sparse subset of ultrasound-sensitive neurons (USSN) (response rate >30%) in V1, comprising only a small fraction of the total population and exhibiting a spatially sparse distribution. The proportion of USSN scale with ultrasound pressure, confirming a direct neuromodulatory effect independent of audition. Intriguingly, despite this sparse activation, ultrasound significantly modulates V1 circuitry: it alters the dynamics of light-sensitive neurons (LSN), with subsets showing excitation, inhibition, or no change in response to visual stimuli. These findings provide the first rigorous *in-vivo* evidence that FUN induces direct mechanical effects on neural activity, disentangling them from auditory confounds. By demonstrating both the specificity and broader circuit-level impact of ultrasound in deafened mice, this study reframes our understanding of FUN’s mechanisms and strengthens its potential as a precise neuromodulatory tool.

**Highlights:**

1. Ultrasound directly activates sparse neurons in deafened mice, demonstrating auditory-independent effects.
2. Sparse ultrasound-sensitive neurons in V1 show pressure-dependent responses.
3. Ultrasound modulates visual circuitry, with diverse excitatory and inhibitory effects.

## Introduction

Focused ultrasound neuromodulation (FUN) integrates non-invasiveness, high spatial resolution, and deep brain penetration, ^1,2^ positioning it as a transformative next-generation tool for neural modulation. ^3,4^ Its potential spans basic neuroscience and therapeutic applications, ^5–7^ with studies demonstrating its ability to elicit behaviors like locomotion in mice, ^8,9^ enhance cognition in non-human primates, ^10^ and improve motion judgment in humans. ^3^ Preclinical models further highlight FUN’s promise for treating disorders such as epilepsy, Alzheimer’s disease, and essential tremor, with clinical successes already reported. ^7,11^ Despite these advances, a critical question looms over the field: does FUN directly modulate neural activity, or are its effects confounded by auditory stimulation?

This uncertainty stems from persistent auditory interference. Sato et al. (2018) showed that ultrasound stimulation of the visual cortex in normal-hearing mice triggered widespread auditory cortex activation, with calcium signal response diminished after chemical deafening. ^12^ Similarly, Guo et al. found that cortical and subcortical responses to ultrasound vanished following auditory nerve transection or cochlear fluid removal, a finding reinforced in genetically deaf mice where low-intensity ultrasound failed to evoke cortical activity. ^13^ These observations suggest that auditory pathways dominate FUN’s apparent effects, casting doubt on its direct mechanical action. Inconsistent motor responses—such as ipsilateral rather than contralateral activation in rodents or unreliable limb movements in larger species—further complicate interpretation. ^14^ Resolving this requires rigorous experiments to isolate ultrasound’s direct neuromodulatory effects from auditory confounds.

Efforts to mitigate auditory interference, such as optimizing pulse waveforms, have shown promise but remain limited. Morteza et al. found that rectangular but not smoothed envelope ultrasound pulses elicited auditory brainstem responses (ABR), suggesting waveform design could reduce auditory artifacts. ^15^ Yet, ABR detects only synchronized activity, potentially missing subtler, asynchronous effects, and device noise may still confound human perception. ^16,17^ Moreover, direct evidence of neuronal activation by smoothed pulses is scarce. We propose that FUN’s effects may be inherently subtler than those of electrical or magnetic stimulation, necessitating sensitive, cellular-resolution tools to detect them in the absence of auditory input.

Current techniques fall short for this task. Functional MRI and ultrasound imaging offer whole-brain views but lack cellular specificity, ^18,19^ while electrophysiology, though direct, is prone to acoustic artifacts and limited in scale. ^20^ Fiber photometry captures specific populations but lack of cellular-level resolution. ^8^ Two-photon calcium imaging (2PCI) addresses these gaps, providing single-cell resolution and broad spatial coverage with minimal invasiveness. ^21^ Recent 2PCI studies have revealed ultrasound’s ability to selectively activate cerebral and cerebellar cortex neurons, yet these studies almost were conducted in normal-hearing animals, leaving auditory confounds unaddressed. ^22–27^

Here, we tackle this challenge using surgically deafened mice and large-scale 2PCI to investigate ultrasound’s direct effects on the V1. We developed a 1.0 MHz ring-shaped transducer compatible with in vivo 2PCI, enabling precise ultrasound delivery in deafened mice. Our findings reveal that ultrasound directly activates a sparse subset of USSN in V1 (response rate >30%), independent of auditory input, and modulates LS neurons responses to visual stimuli. These results provide definitive evidence of FUN’s mechanical neuromodulatory capacity, refining its mechanisms and reinforcing its potential as a precise tool for neural circuit manipulation.

## Results

### Experimental setup and data processing

We engineered an annular ultrasound transducer integrated with a 16× objective lens (focal length 3 mm), positioning the transducer surface coplanar with the lens’s lower edge to maintain a fixed ultrasound-imaging plane alignment (Figure 1A). The acoustic field, measured via hydrophone post-cranial window penetration, exhibited a well-defined spatial profile, with normalized distributions mapped in longitudinal (XZ) and transverse (XY) planes (Figure 1B). For stimulation, we employed smoothed ultrasonic pulses (burst duration 0.5 s, stimulation interval 30 s, 50% duty cycle; Figure 1C), designed to minimize auditory artifacts.

**Figure 1.**
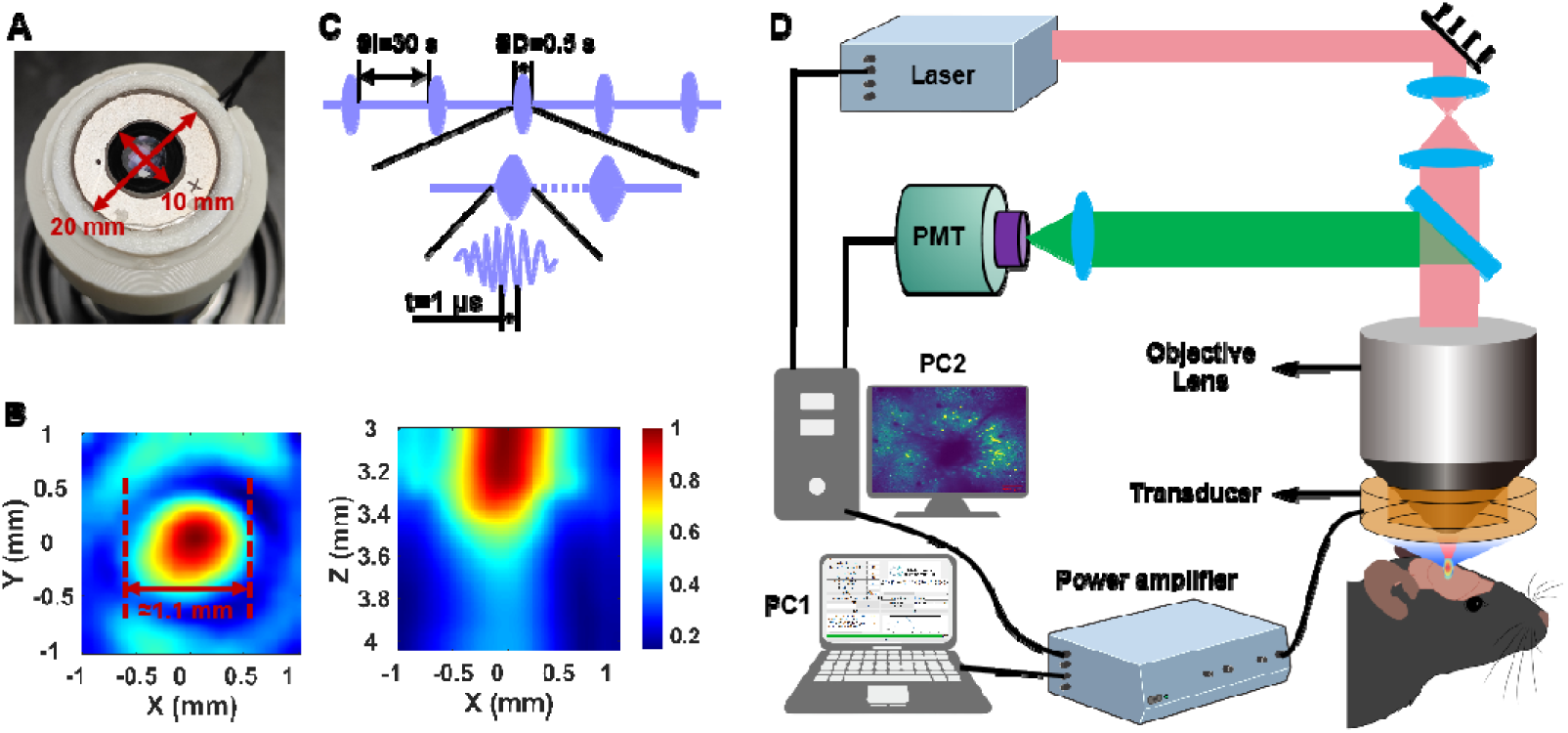
Schematic diagram of the experimental system. (A) Photograph of the custom-designed annular ultrasound transducer integrated with the objective lens. (B) Normalized ultrasound field distribution in the XY and XZ planes generated by the transducer after passing through the cranial window. (C) Schematic representation of the ultrasound stimulation waveform and parameters. Diagram illustrating the integration of the 2PCI system with the ultrasound neuromodulation setup.

The experimental system synchronized two-photon calcium imaging with ultrasound delivery (Figure 1D). A personal computer (PC1) controlled the ultrasonic drive system to generate precise waveforms to drive the transducer, while sending synchronous triggers to PC2. Then, PC2 running ScanImage ^28^ to directe a femtosecond laser to scan the cranial window, exciting GCaMP6s in V1 neurons of surgically deafened mice. A photomultiplier tube (PMT) captured emitted signals, reconstructing time-series datasets of neuronal GCaMP6s fluorescence intensity (ΔF/F). Imaging parameters included a 0.75 × 0.75 mm field of view (512 × 512 pixels) at a frame rate of 30 Hz, ensuring high-resolution monitoring of large neuronal populations. This setup enabled precise assessment of ultrasound’s direct effects on V1, free from auditory confounds, laying the groundwork for subsequent analyses of neuronal responses.

### Global calcium signal responses in V1 induced by ultrasound and visual stimuli

To eliminate auditory confounds, we performed cochlear removal in mice expressing rAAV-hSyn-GCaMP6s in V1. We confirmed that the auditory of the mice was deprived by comparing the ABR signals of mice before and after cochlear removal. via absent ABR (Figure 2A, B). We first tested V1 responsiveness using pulsed blue light (50 ms pulse width, 500 ms duration, 10 Hz PRF) delivered to the eyes, paired with two-photon calcium imaging of contralateral V1 (Figure 2C). Visual stimulation robustly activated neurons (yellow arrow, Figure 2C), with average GCaMP6s fluorescence (ΔF/F) across the 0.75 × 0.75 mm field rising significantly within 3 s, followed by suppression from 5–8 s. This biphasic response confirms intact V1 function in deafened mice.

**Figure 2.**
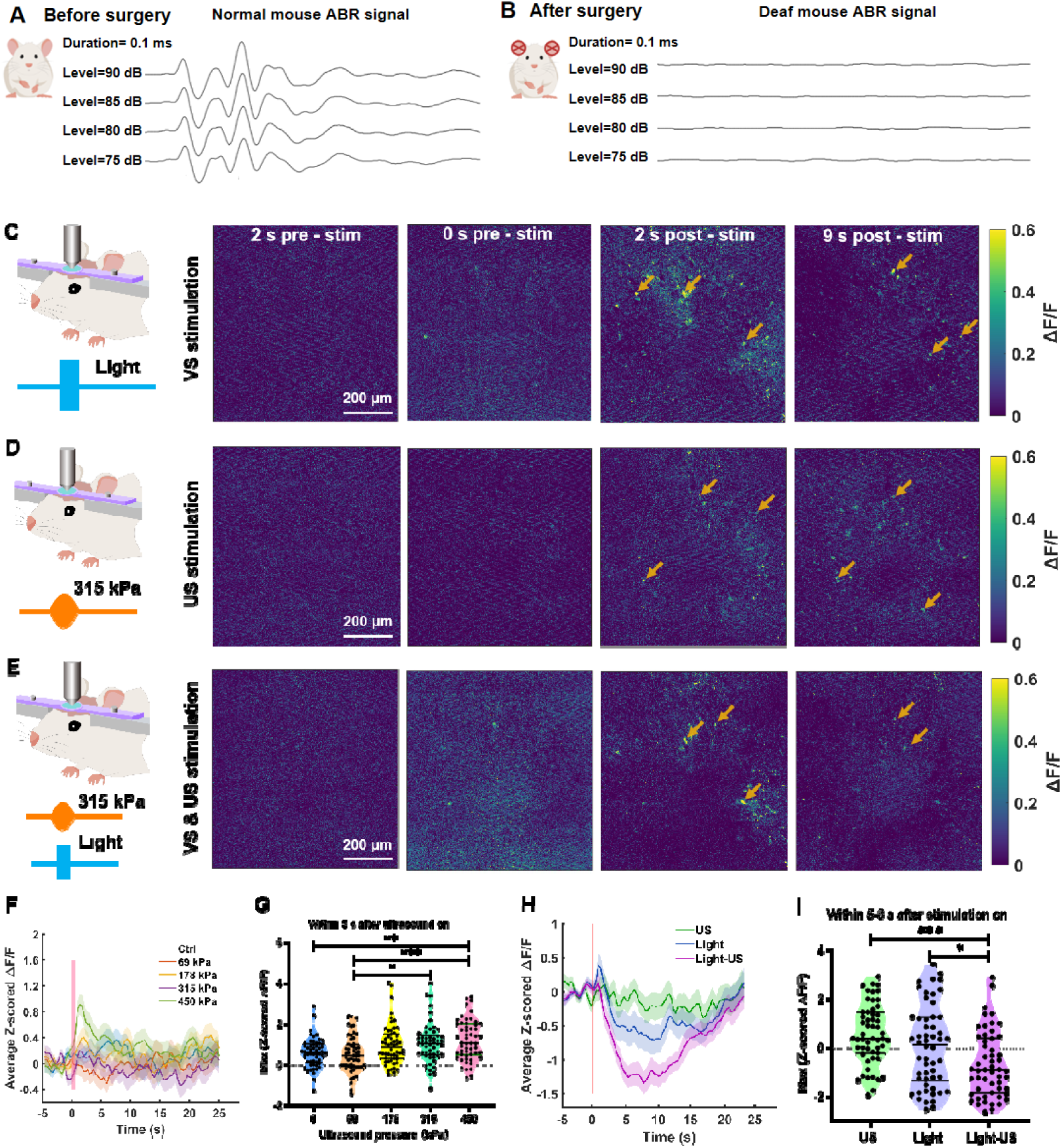
Ultrasound directly activates neurons in the V1 of deaf mice. (A, B) Representative ABR signals from a wild-type mouse before and after surgical deafening. (C) ΔF/F traces from a representative 2PCI field before and after visual stimulation (blue light, pulse width: 50 ms, stimulation duration: 500 ms, PRF: 10 Hz). (D) ΔF/F traces from a representative 2PCI field before and after ultrasound stimulation (center frequency: 1 MHz, negative peak pressure: 315 kPa, pulse width: 500 ms, stimulation duration: 50 ms, PRF: 10 Hz). (E) ΔF/F traces from a representative 2PCI field before and after simultaneous visual and ultrasound stimulation. (F) Average z-scored ΔF/F in the V1 of mice under ultrasound stimulation at a PRF of 10 Hz with varying pressures (n = 6 mice). (G) Maximum z-scored ΔF/F within 3 seconds after ultrasound stimulation at different pressures (negative peak pressure: 0–450 kPa, pulse width: 50 ms, center frequency: 1 MHz, stimulation duration: 500 ms, PRF: 10 Hz) (n = 6 mice; 9 trials per group; bars represent mean ± SEM; *P < 0.05, **P < 0.01, ***P < 0.001, by one-way ANOVA followed by Sidak’s post-hoc multiple comparison test). (H) Average z-scored ΔF/F across the entire 2PCI field of view in the V1 of mice under visual stimulation, ultrasound stimulation (315 kPa), and simultaneous visual and ultrasound stimulation (315 kPa). (I) Maximum z-scored ΔF/F measured 5–8 seconds after different stimulation conditions (n = 6 mice; 9 trials per group; bars represent mean ± SEM; *P < 0.05, **P < 0.01, ***P < 0.001, one-way ANOVA followed by Sidak’s post-hoc multiple comparison test).

We then applied ultrasound to V1 at pressures of 0, 69, 178, 315, and 450 kPa (500 ms pulse width, 1 MHz, 50 ms duration, 10 Hz PRF, 30 s interstimulus interva). At pressures below 450 kPa, ultrasound elicited no significant change in average ΔF/F across the imaging field (Figure 2F). At 450 kPa, however, ultrasound markedly increased V1 calcium signals (Figure 2F, G). Notably, at 315 kPa, a subset of neurons showed activation (Figure 2D), hinting at selective modulation. To test this, we combined 315 kPa ultrasound with visual stimulation (10 Hz PRF, 500 ms pulse width, 50 ms duration, and 30 s interstimulus interval) (Figure 2E). Ultrasound enhanced the suppression phase of the visual-evoked calcium response, reducing overall ΔF/F (Figure 2H, I). These field-wide averages suggest ultrasound modulates V1 activity, with effects varying by pressure and stimulus context. Single-neuron analyses follow to dissect these dynamics further.

### Ultrasound directly activates sparse USSN in deafened mice

To characterize ultrasound’s direct effects on V1, we analyzed 2PCI datasets from deafened mice (Figure 3A, B). Representative GCaMP6s traces (z-scored ΔF/F) revealed heterogeneous neuronal responses to 450 kPa ultrasound stimulation (10 Hz PRF): some neurons fired frequently, others sporadically, and some only post-stimulation (Figure 3B). We developed a 2PCI pipeline to quantify these dynamics (Figure 3C): time-series images underwent motion correction, followed by semi-automated ROI segmentation (manually refined) to identify neurons. Fluorescence signals were extracted, aligned to ultrasound timing (-5 to +25 s), and ΔF/F calculated. Neurons responding within 3 s post-stimulation were classified as ultrasound-sensitive (USS, response rate >30%) or non-ultrasound-sensitive (NUSS, 0 < response rate ≤ 30%), with spatial coordinates mapped.

**Figure 3.**
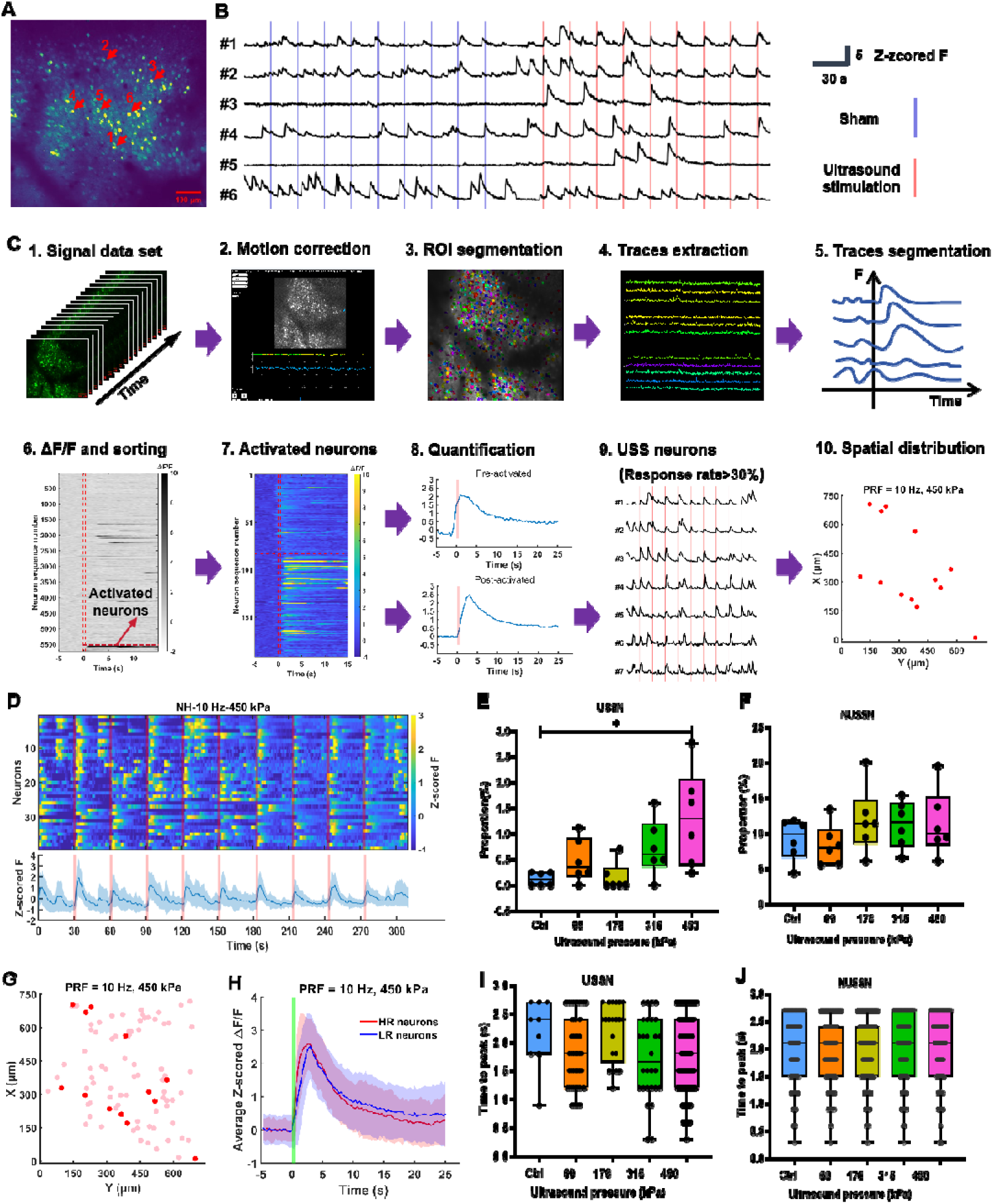
Ultrasound directly activates sparse USSN in the V1 of deaf mice. (A) Representative projections of 2PCI datasets. Representative 2PCI projection images.(up to you to use the world dataset or image) (B) Representative z-scored ΔF/F traces of neurons without and with ultrasound stimulation. (C) Schematic workflow for extracting and processing GCaMP6s fluorescence signals from individual neurons. (D) Heatmap and mean traces of z-scored ΔF/F for USSN activated by ultrasound stimulation at a pulse repetition frequency (PRF) of 10 Hz and varying pressures (69, 178, 315, and 450 kPa) (from 2,693 neurons of 6 mice). (E, F) Proportion of USSN and Non-USSN under 10 Hz ultrasound stimulation at different pressures (n = 6 mice with 2,693 neurons per group; bars represent mean ± SEM; *P < 0.05, one-way ANOVA followed by Sidak’s post-hoc multiple comparison test). (G) Spatial distribution of representative USSN (dark red circles) and Non-USSN (light red circles) under 450 kPa ultrasound stimulation. (H) Averaged z-scored ΔF/F traces of USSN (red traces) and Non-USSN (blue traces) under 450 kPa ultrasound stimulation. (I, J) Average time for z-scored ΔF/F traces of USSN and Non-USSN to reach peak values under different ultrasound pressures (n = 6 mice, 2,693 neurons per group; bars represent mean ± SEM; *P < 0.05, one-way ANOVA followed by Sidak’s post-hoc multiple comparison test).

Across 2,693 neurons from six mice, USSN comprised a sparse subset (Figure 3D). At 450 kPa, only 1.28% were USSN, despite a pressure-dependent increase above 178 kPa (Figure 3E). NUSSN, conversely, peaked at 178 kPa without further escalation (Figure 3F). Spatially, USSN (dark red) appeared randomly dispersed amid NUSSN (light red) under 450 kPa (Figure 3G). USSN exhibited steeper ΔF/F slopes (red traces) than NUSS (blue traces; Figure 3H), with shorter time-to-peak as pressure rose, a trend absent in NUSSN (Figure 3I, J). These data confirm ultrasound directly activates a sparse, pressure-sensitive USS population, while NUSSN responses may reflect spontaneous or indirect effects, highlighting FUN’s selective neuromodulatory capacity.

### Ultrasound alters LS neuron responses to visual stimulation

We screened V1 neurons for light-sensitivity (LS) and ultrasound-sensitivity (USS) using visual stimulation, 315 kPa ultrasound, and combined conditions, analyzing calcium signals via 2PCI (as above). Spatial maps (Figure 4A–C) show USSN/LSN (Dark circles) sparsely distributed among non-sensitive (NUSS/NLS) neurons (light circles) under different stimulation conditions. Proportions of USS, LS, and dual-sensitive (LS-USS) neurons varied by condition (Figure 4D), with ultrasound boosting LS neuron prevalence during visual stimulation. Heatmaps (Figure 4E) and average traces (Figure 4F, G) revealed distinct responses: visual stimulation alone drove LS neurons but not USS, while ultrasound alone activated USSN with minimal LS impact (Figure 4H). Venn analysis confirmed minimal overlap between LS and USS populations (Figure 4I), indicating separate cohorts responsive to each modality.

**Figure 4.**
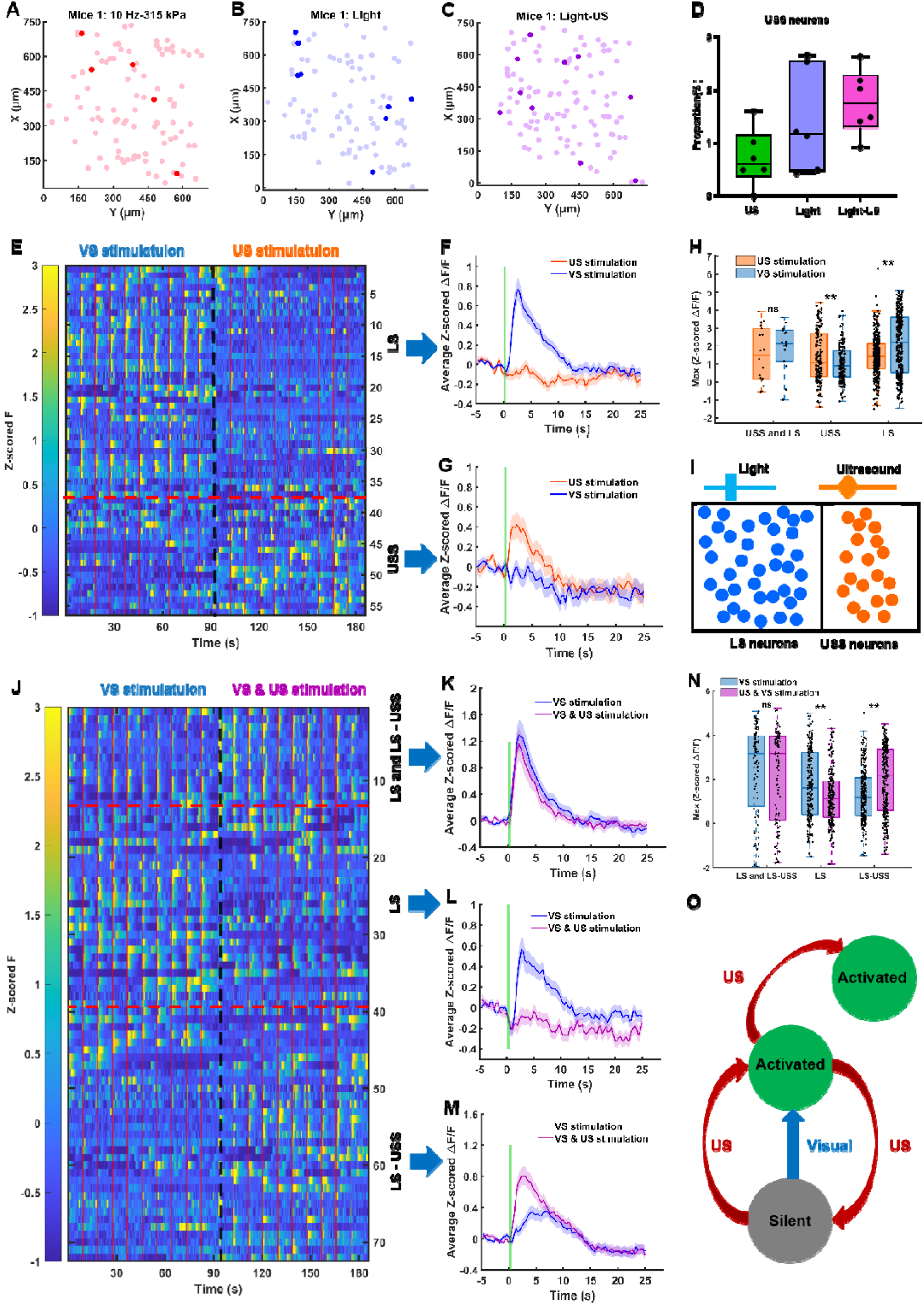
Ultrasound modulates the response of LSN in V1 to visual stimulation. (A) Spatial distribution of USSN (dark red circles) and Non-USSN (light red circles) neurons under 315 kPa ultrasound stimulation. (B) Spatial distribution of LSN (dark blue circles) and Non-LSN (light blue circles) neurons under visual stimulation. (C) Spatial distribution of LSN+USSN (dark purple circles) and other neurons (light purple circles) neurons under simultaneous ultrasound and visual stimulation. (D) Proportion of USSN under different stimulation conditions (n = 6 mice, 2,693 neurons per group; bars represent mean ± SEM; *P < 0.05, one-way ANOVA followed by Sidak’s post-hoc multiple comparison test). (E) Heatmap of GCaMP6s fluorescence intensity (ΔF/F) for LSN and USSN under visual and ultrasound stimulation (n = 6 mice, 2,693 neurons per group). (F, G) Average ΔF/F traces of LSN and USSN in response to visual and ultrasound stimulation. (H) Maximum ΔF/F values of LSN+USSN, LSN, USSN) within 3 seconds after different stimulation conditions (each stimulation repeated 9 times; bars represent mean ± SEM; two-tailed t-test, *P < 0.05, **P < 0.01). (I) Correlation between the populations of LSN and USSN. (J) Heatmap of z-scored ΔF/F for LSN and USSN under simultaneous visual and ultrasound stimulation. (K-M) Average z-scored ΔF/F traces of LSN+USSN, LSN, USSN in response to visual stimulation and simultaneous ultrasound and visual stimulation. (N) Maximum z-scored ΔF/F values of LSN+USSN, LSN, USSN within 3 seconds after different stimulation conditions (each stimulation repeated 9 times; bars represent mean ± SEM; two-tailed t-test, *P < 0.05, **P < 0.01). Schematic representation of the excitatory and inhibitory effect of ultrasound on LSN in response to visual stimulation.

Under combined stimulation, ultrasound modulated LS responses (Figure 4J). Subsets of LS and LS-USSN showed unchanged visual responses (Figure 4K, N), but ultrasound inhibited a distinct LS subset (Figure 4L, N) while enhancing some non-light-sensitive (NLS) neurons’ visual responses (Figure 4M, N). Summary data (Figure 4O) demonstrates ultrasound’s capacity to bidirectionally regulate V1 circuitry, with sparse USSN directly activated and LS neurons diversely modulated—some inhibited, others enhanced—under task-relevant conditions, offering a framework for dissecting FUN’s network-level effects.

## Discussion

Our findings establish that ultrasound directly modulates V1 activity in deafened mice, independent of auditory interference. Below 450 kPa, no field-wide calcium response emerged, yet at 315 kPa, sparse ultrasound-sensitive (USS) neurons (0.72% of total) were activated, rising to 1.28% at 450 kPa. Combining 315 kPa ultrasound with visual stimulation revealed network-level modulation-enhancing visual-evoked suppression macroscopically while diversely altering LS neuron responses (inhibition, enhancement, or no change). These results demonstrate that low-intensity ultrasound exerts subtle, task-like effects ^29^ rather than broad excitation or inhibition, challenging its conflation with auditory artifacts and affirming its neuromodulatory potential.

Why does ultrasound reliably evoke limb movement in mice but not larger species? In deafened mice, sparse USS activation (1% at 450 kPa) scaled with pressure, yet remained modest—potentially too weak to drive motor output in larger cortices with lower ultrasound penetration efficiency. Mice’s broader auditory range (1–100 kHz) versus primates (1–60 kHz) or humans (0.02–20 kHz) may also amplify indirect effects in hearing models, misattributing motor responses to ultrasound. Our findings align with Sato et al. ^12^ and Guo et al, ^13^ where low-pressure ultrasound yielded no direct activation in deafened or nerve-sectioned animals, reinforcing that FUN’s direct impact is subtle and localized.

Ultrasound’s modulation of V1 holds promise for neuroscience and therapy. Beyond dissecting visual cortex dynamics—key to perception, plasticity, and multisensory integration ^32,33^—it could inspire vision restoration strategies, as seen with illusory percepts in prior studies. Unlike TMS or tDCS, which robustly drive motor responses, ^34–36^ FUN’s subtlety suggests applications in nuanced circuit manipulation, such as brain-computer interfaces or psychiatric treatments targeting aberrant V1 activity (e.g., depression, migraine). ^32,37^ However, USSN’ identity remains unclear. Our previous investigations and emerging evidence collectively demonstrate that the mechanosensitive ion channel Piezo1 serves as a critical mediator in ultrasound-mediated neuromodulation, exhibiting a sparse expression pattern in brain that may similar to the distribution characteristics of ultrasound-sensitive (USS) neurons. ^27,30,31^ Notably, recent studies have reported that TRPM2 and TRPC6 channels may also play significant roles in ultrasound neuromodulation. ^9,39^ However, these studies may have been subject to auditory interference. Future work should map their morphology, probe mechanosensitive channels via sequencing, ^38^ and test diverse cell types across regions. Optimized parameters, free of auditory bias, will further refine FUN’s precision.

In sum, ultrasound directly activates sparse USSN and subtly tunes V1 networks, mimicking task-related states. This reframes FUN as a precise, mechanically driven tool with transformative potential for research and clinical intervention.

## Methods

### Ultrasound field simulation

The acoustic field of an annular ultrasonic transducer (inner diameter: 11 mm, outer diameter: 19 mm) was simulated using the k-Wave MATLAB toolbox. ^40^ Spatial and temporal grid steps were 200 μm and 5 ns (CFL = 0.1 in water), with a 200 MHz sampling rate. Free-field water parameters were density 1000 kg/m³, sound speed 1500 m/s, and attenuation 0.002 dB/(MHz²·cm). For transcranial simulations, a 0.6-mm-thick, 6-mm-wide glass cranial window was modeled (density 2400 kg/m³, sound speed 5600 m/s, attenuation 0.02 dB/(MHz²·cm)), with air outside and the transducer 2.0 mm from the window surface.

### Fabrication and testing of the annular ultrasonic transducer

A 3D-printed housing (inner diameter: 20.5 mm, outer diameter: 23 mm) encased a PZT-4 piezoelectric ceramic. Electrodes were bonded with conductive silver adhesive (3022KIT, Von Roll USA), cured for 12 h. The ceramic was secured in the housing with epoxy resin (AB adhesive) mixed with 25% aluminum oxide powder, with thin layers applied to both surfaces for insulation. After 24 h curing, the assembly was bonded to a 3D-printed holder nested with the objective lens. Acoustic fields in water and through a 0.6-mm cranial window (from post-mortem mouse skulls) were measured 2.0 mm from the transducer using a calibrated hydrophone and lab-built scanning system, ^41^ mapping transverse and longitudinal profiles.

### Animal preparation and virus injection

Six male C57BL/6 mice (8 weeks, 20–25 g) were housed under standard conditions (12-h light/dark cycle, 22 ± 1°C, ad libitum food/water). Procedures were approved by the Guangdong Institute of Intelligence Science and Technology’s IACUC, adhering to ethical guidelines. Mice were anesthetized with ketamine (10 mg/mL) and xylazine (2 mg/mL) i.p., fixed stereotaxically, and craniotomized at AP -2.8 mm, ML -2.4 mm from bregma. rAAV-hSyn-GCaMP6s was injected into V1 (DV -1.0 mm) via microinjection pump and glass micropipette. After 10 min, the scalp was sutured, cefazolin administered i.p., and mice recovered for 2 weeks for viral expression.

### Transparent cranial window implantation

Custom glass plugs (4-mm and 6-mm discs) were bonded with transparent epoxy. Under anesthesia (ketamine/xylazine i.p.), a 5-mm skull section over the injection site was drilled, preserving the dura. The plug was sealed with dental cement, and a head-fixation apparatus was cemented to the skull for imaging stability. ^42^

### Deafness surgery

Anesthetized mice (ketamine/xylazine i.p.) underwent cochlear ablation. The ear canal was opened, tympanic membrane and ossicles removed, cochlea excised, and fluid drained to disrupt audition. The site was sutured, and post-operative care included pain management.

### 2PCI with ultrasound and visual stimulation

Mice were head-fixed on a motorized stage, acclimated over multiple sessions. The transducer, nested with a 16× objective, was stabilized with parafilm and coupled to the cranial window via ultrasound gel. The 1 MHz ultrasound system (41) delivered customizable pulses (PRF: 10, 1000, 2000 Hz; pressures: 69, 178, 315, 450 kPa), synchronized with 2PCI (512 × 512 pixels, 3 Hz, 0.75 × 0.75 mm FOV). Nine of ten stimulations per trial were analyzed (first discarded). Visual stimulation used a blue LED (0.5 s, 1 cm from left eye), isolated or paired with ultrasound, with synchronization split to trigger 2PCI and LED.

### 2PCI data processing

Suite2p preprocessed GCaMP6s data: motion correction, ROI segmentation (manually curated), and signal extraction (ROI minus background). ^43^ Epochs spanned -5 to +25 s around stimuli, with ΔF/F and z-scores calculated. Excited neurons (z-scored ΔF/F > 2 within 3 s post-stimulus) were identified; pre-stimulated active neurons were excluded. Ultrasound- or light-sensitive neurons had response rates >30%.

## Acknowledgments

This work was supported in part by the National Natural Science Foundation of China (32371151), Guangdong High Level Innovation Research Institute (2021B0909050004), the Hong Kong Research Grants Council Collaborative Research Fund (C5053-22GF), General Research Fund (15224323 and 15104520), Hong Kong Innovation Technology Fund (MHP/014/19), internal funding from the Hong Kong Polytechnic University (G-SACD and 1-CDJM).

## Author contributions

**Conceptualization:** ZHQ, JRH, JJZ, **Methodology:** JRH, JJZ, XXW, ZHC, JY, ZY. **Funding acquisition:** ZHQ, LS. **Supervision:** ZHQ. **Competing interests:** The authors declare that they have no competing interests. **Data and materials availability:** All data are available in the main text or the supplementary materials.

